# A supervised learning framework for chromatin loop detection in genome-wide contact maps

**DOI:** 10.1101/739698

**Authors:** Tarik J. Salameh, Xiaotao Wang, Fan Song, Bo Zhang, Sage M. Wright, Chachrit Khunsriraksakul, Feng Yue

## Abstract

Accurately predicting chromatin loops from genome-wide interaction matrices such as Hi-C data is critical to deepen our understanding of proper gene regulation events. Current approaches are mainly focused on searching for statistically enriched dots on a genome-wide map. However, given the availability of a wide variety of orthogonal data types such as ChIA-PET, GAM, SPRITE, and high-throughput imaging, a supervised learning approach could facilitate the discovery of a comprehensive set of chromatin interactions. Here we present Peakachu, a Random Forest classification framework that predicts chromatin loops from genome-wide contact maps. Compared with current enrichment-based approaches, Peakachu identified more meaningful short-range interactions. We show that our models perform well in different platforms such as Hi-C, Micro-C, and DNA SPRITE, across different sequencing depths, and across different species. We applied this framework to systematically predict chromatin loops in 56 Hi-C datasets, and the results are available at the 3D Genome Browser (www.3dgenome.org).

## INTRODUCTION

The proper gene regulatory program of mammalian cells are largely influenced by the 3D conformation of chromosomes^1^. At kilobase to megabase scales, gene promoters are often connected to their distal regulatory elements, such as enhancers, through chromatin loops; rewiring of such loops has been implicated in developmental diseases and tumorigenesis^2,3^. It has been shown that chromatin loops are mediated by architectural proteins CTCF and cohesin via a loop extrusion model, where CTCF binds to a specific and non-palindromic motif in a “convergent” orientation at two sites, acting as loop anchors^4,5^.

A growing number of experiments have been used to detect chromatin loops. Hi-C^6^, a high-throughput derivative of Chromosome Conformation Capture (3C)^7^, quantifies contacts between all possible pairs of genomic loci using a proximity-ligation procedure. With an improved experimental protocol and deep sequencing, *in-situ* Hi-C^8^ makes it possible to detect loops at kilobases. By introducing micrococcal nuclease for chromatin fragmentation instead of restriction enzymes, Micro-C^9^ further enables nucleosome-resolution analysis of chromatin interactions. Proximity-ligation techniques also include ChIA-PET^10^, PLAC-Seq^11^, and HiChIP^12^, which detect loops bound to target proteins through chromatin immunoprecipitation steps, and include Capture C^13^ and Capture Hi-C^14^, which enrich interactions with a given set of sequences. Recently, several ligation-free techniques emerged to measure different aspects of chromatin organization. Genome Architecture Mapping (GAM)^15^ quantifies chromatin contacts by sequencing DNA from a set of ultrathin nuclear sections at random orientations. Trac-looping^16^ captures multiscale contacts by inserting a transposon linker between interacting regions. DNA SPRITE^17^ follows a split-pool procedure to assign unique barcodes to individual complexes, with read pairs sharing identical barcodes treated similarly to contacts in Hi-C. Besides these biomedical protocols, high-throughput imaging approaches such as STORM^18^ and HiFISH^19^ can directly measure spatial distances at the single-cell level.

Several computational tools have been developed to identify chromatin loops accordingly. For Hi-C data: Fit-Hi-C^20^ performs a distance-dependent spline fitting procedure to refine its global background and chooses binomial distribution as the null model to evaluate contact significance, which can output ~1 million cis-interactions with deeply sequenced reads^21^. HiCCUPS^8,22^ incorporates local background into its model and utilizes the Poisson test with a modified Benjamini-Hochberg adjustment to determine statistical significance, and generally reports thousands of loop interactions. Analysis of ChIA-PET data always starts with peak calling to identify anchor regions of the target protein, however the background assumption used in interaction identification varies among tools. For example, the first published ChIA-PET software tool^23,24^ adopts a hyper-geometric distribution to filter out noise, but the more recent Mango^25^ builds a null model by incorporating both the genomic distance and read depth of each anchor. For PLAC-Seq and HiChIP, the recently developed MAPS^26^ filters original interactions against ChIP-Seq peaks of the same protein, and conducts a specific normalization procedure before evaluating significance. We observe that nearly all available tools are based on testing for significant enrichment compared to a local or global background, with the specific calculations being quite empirical and difficult to generalize between techniques. It would be intriguing and potentially beneficial to automatically distinguish loop vs non-loop interactions in a data-driven manner, which is a standard supervised learning task.

Machine learning (ML) has been successfully applied in genomics settings, such as predicting microRNA target activities^27^, annotating chromatin states^28,29^, and characterizing functional effects of noncoding variants^30^. In chromatin conformation studies, manifold learning strategies are employed by miniMDS^31^ and GEM^32^ to estimate 3D structures from 2D contact maps. Recently, efforts have been made to predict 3D interactions from 1D sequence and epigenomics datasets^33,34^. In addition, we developed HiCPlus software^35^, which can greatly enhance the Hi-C data resolution through a deep convolutional neural network. So far, the potential benefits of ML approaches for loop detection at kilobase scales are relatively unexplored.

Here we present Peakachu (**U**nveil **H**i-**C A**nchors and **Peak**s) (Fig. 1), a supervised ML framework to detect chromatin loops on genome-wide interaction maps. Peakachu builds loop-classifying models from defined positive and negative training sets: the positive set could be any kind of interactions from either biologically enriched experiments such as ChIA-PET and HiChIP, or a high-throughput imaging experiment such as HiFISH. The negative set is generated from loci randomly sampled from two populations: 1) contacts with genomic distances similar to the positive set and 2) contacts with larger genomic distances than the positive set. Once the training set is defined, Peakachu applies a hyperparameter search to find the best random forest model separating the two classes, which can be used to detect loops in genome-wide contact maps. We show that the predictions made by Peakachu have high precision and recall rates. Further, we demonstrate Peakachu can detect high-resolution chromatin loops with only 200-300 million reads. With pre-trained models, we successfully predict chromatin loops in 56 Hi-C datasets with different sequencing depths and make them available at the 3D Genome Browser (3dgenome.org). Finally, we show Peakachu is a platform-agnostic tool by applying it in two additional platforms, Micro-C and DNA SPRITE.

**Figure 1.**
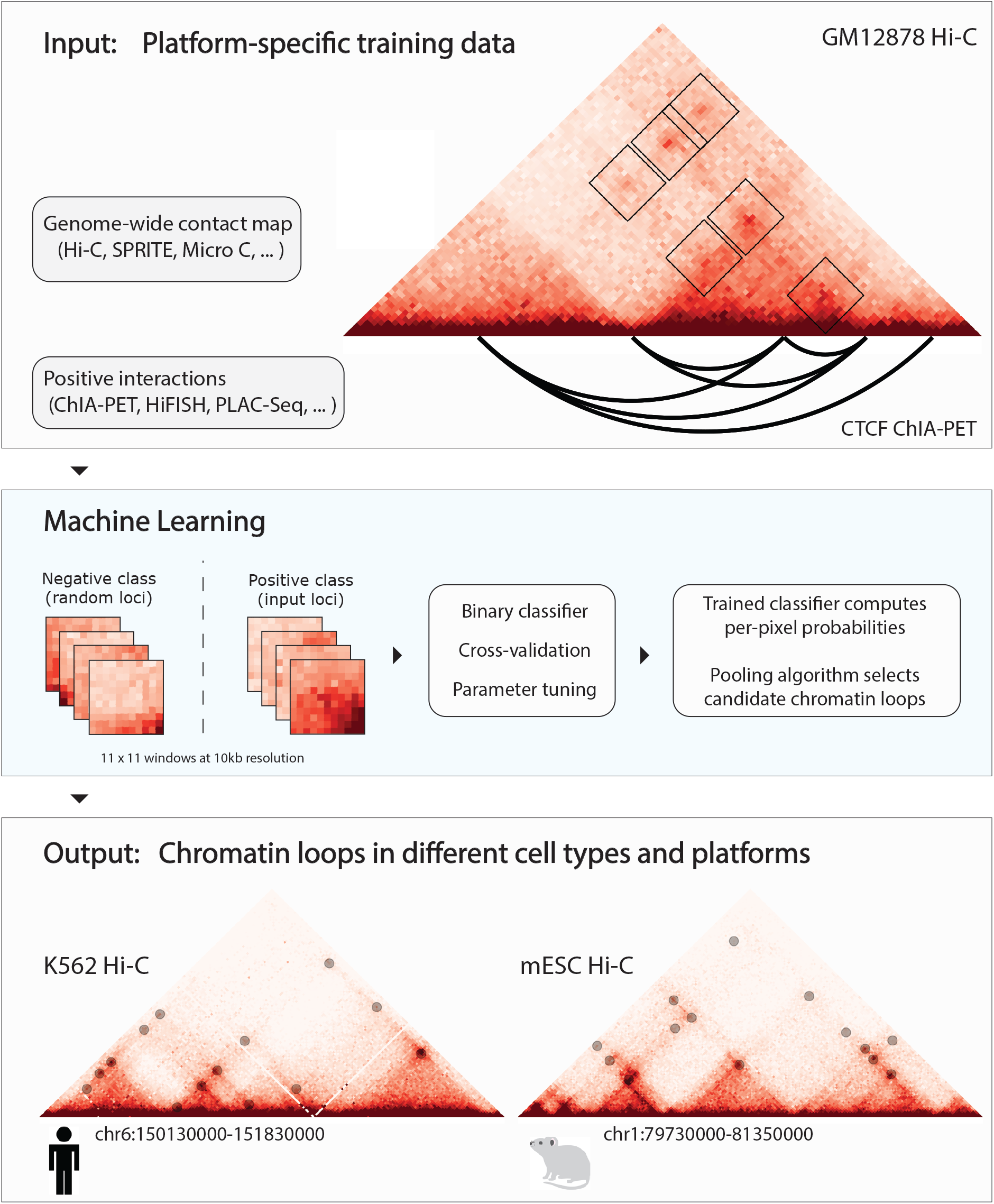
A binary classification framework for loop detection in genome-wide contact data. A contact matrix from Hi-C or similar experiment is decomposed into a training set defined by subwindows either centered at positive interactions from an orthogonal method (ChIA-PET, PLAC-Seq, HiFISH, …) or random loci of similar genomic distance. Hyperparameter tuning within a 3-fold cross-validation is applied to select a random forest model best able to distinguish the two classes. A trained model can then compute per-pixel probabilities in a different contact map from the same platform, with Hi-C depicted here. A greedy pooling algorithm selects the best-scored contacts from clusters of high probability pixels.

## Results

### Overview of the Peakachu framework

We describe the overall approach by Peakachu in Figure 1. There are two parts of the input. The first part is a genome-wide interaction matrix, such as Hi-C or Micro-C data. The second part consists of the positive and negative training datasets. Positive training sets are defined by loops identified from orthogonal techniques such as ChIA-PET, PLAC-Seq, Capture Hi-C, or even high-resolution imaging data as they become available. For negative training sets, an equal number of pixels are randomly selected from nonzero values from a distance distribution derived from the positive set. The negative set always contains contacts with the similar genomic distance resembling the positive set, and a set of long-range contacts resembling the noise inherent in contact maps.

The feature vectors of training samples are defined by the surrounding pixels of each sample. Each vector includes the absolute value of each pixel as well as the relative rank of each pixel within the sample. The exact window size is configurable, while we used 11×11 windows at 10kb resolution for all work presented in this text. Once the positive and negative sets are defined as feature vectors, a 3-fold cross-validation procedure is applied to train a random forest model best separating the two classes. Briefly, the input is randomly separated into three equal parts and multiple models are trained using several combinations of tuning parameters. Each of these models is trained on two parts of the data and one part is used for scoring. The parameter combination achieving the best score is selected to fit a final model using the whole training set.

In the prediction stage, similar feature vectors are defined for all nonzero values in a contact map to compute per-pixel probability scores, and a pooling algorithm is applied to eliminate local loop redundancy. Note the model trained on one dataset can be used to predict loops on other matrices from the same platform. As shown in the following sections, a model trained on one cell type can be used to predict loops from Hi-C matrices in other cell types with comparable performance. Detailed description of the framework can be found in the methods section.

### Different orthogonal datasets reveal different sets of chromatin interactions

To train and evaluate the performance of Peakachu, we decided to use the high resolution Hi-C data in lymphoblast cell line GM12878^8^, a tier one ENCODE cell line with extensive epigenome data available. There are four types of orthogonal data in this cell line: CTCF ChIA-PET^23^, RAD21 ChIA-PET^36^, SMC1 HiChIP^12^, and H3K27ac HiChIP^37^ (Supplementary Table 1). First, we observed that each of these four ChIP-based assays predicted a unique set of chromatin interactions (Supplementary Fig. 1). Among them, CTCF ChIA-PET identified the highest number of chromatin loops, while H3K27ac HiChIP only identified 6,395 loops. 39% (2,499 out of the 6,395) H3K27ac HiChIP loops were predicted by all four techniques, while 22% (1,432/6,395) are uniquely predicted in this dataset. Similarly, 49% of the CTCF ChIA-PET loops are unique, while only 13% of them can be recovered by all techniques, potentially due to the fact that the number of loops in CTCF ChIA-PET is much larger than other data types.

More interestingly, we found that these four datasets identified chromatin loops at different genomic distance. For example, 75% (4,810/6,395) of the H3K27ac HiChIP loops are within 250kb (Fig. 2a), and only 8% (500/6,395) are over 500kb. On the contrary, 42% (23,420/55,222) of the pooled CTCF ChIA-PET interactions are long-range (>500kb) and only 34% are short-range (<250kb). This is consistent with the previous observation that CTCF is more responsible for long-range interactions^38,39^, and it has been shown to be a key component of the loop extrusion model^4,5^ and the formation of TADs^40^. At the same time, H3K27ac is a histone mark for active enhancers and promoters, and therefore it is possible the contacts enriched for H3K27ac are shorter-range interactions between promoters and enhancers, which could be more dynamic than CTCF loops^41^. Therefore, in the following sections, we will first evaluate the models trained by both CTCF ChIA-PET and H3K27ac HiChIP and then combined their results to achieve a more comprehensive set of predictions from Hi-C data.

**Figure 2.**
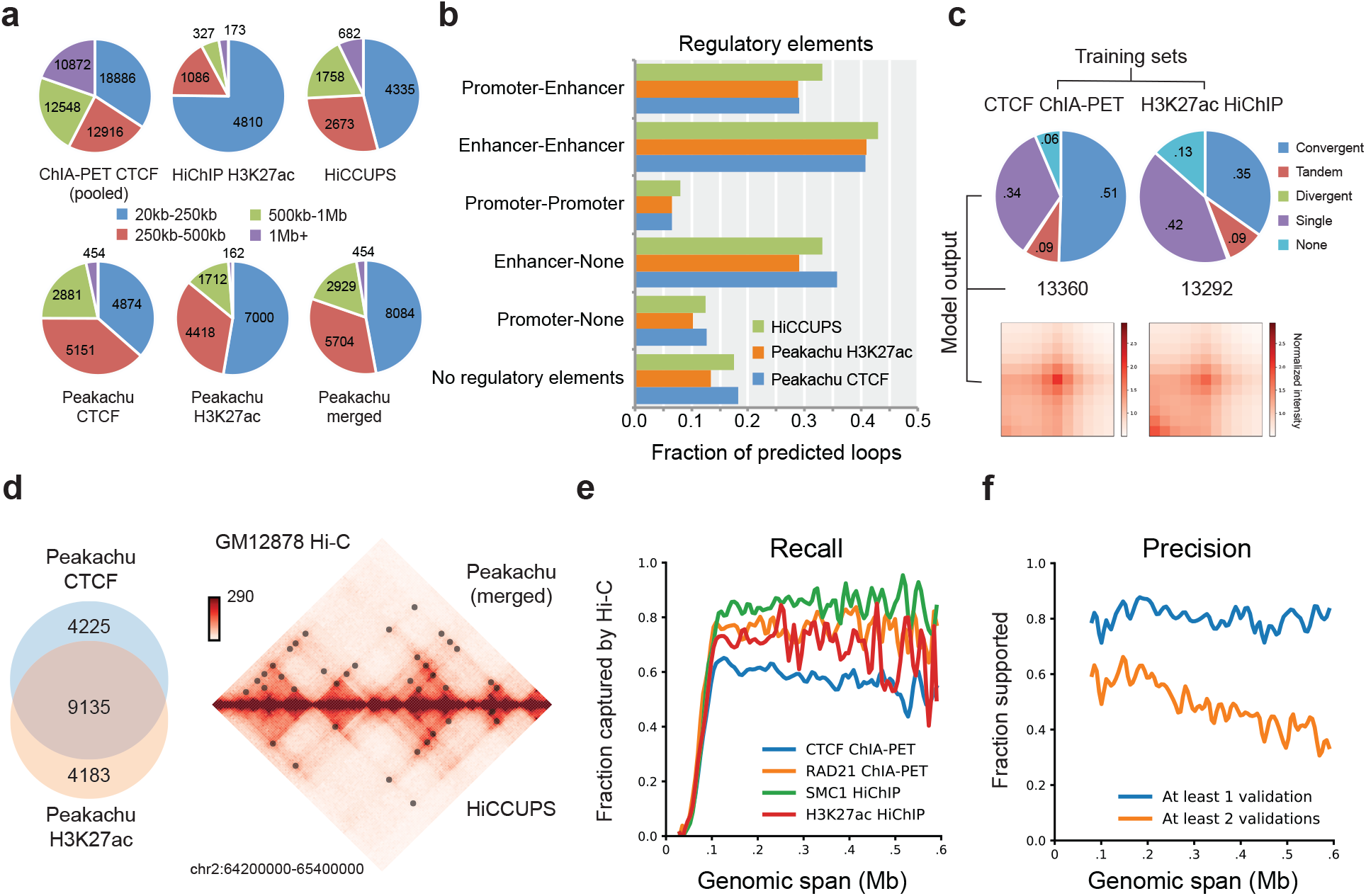
Peakachu framework applied in GM12878 Hi-C. **a.** Distance distributions of CTCF ChIA-PET, H3K27ac HiChIP, and HiCCUPS (Hi-C) interactions in GM12878 (top row). Distributions of Peakachu loops predicted from Hi-C after training with CTCF ChIA-PET or H3K27ac HiChIP data, and union of both predictions (bottom row). Interactions in CTCF ChIA-PET were first pooled to remove local redundancy with the same algorithm used by Peakachu. **b.** Proportion of predicted loops with different regulatory element combinations at anchor loci. **c**. CTCF binding patterns and APA analysis of Peakachu predictions. **d**. Overlap of loops predicted by Peakachu models trained with either CTCF ChIA-PET or H3K27ac HiChIP examples, and visualization of interactions predicted from Hi-C of GM12878 by Peakachu and HiCCUPS. **e**. Fraction of GM12878 interactions in orthogonal experiments recaptured by merged Peakachu loops. **f**. Fraction of Peakachu predictions validated by orthogonal experiments.

### Peakachu accurately captures known interactions from Hi-C data

We first trained our model in GM12878 Hi-C data with the 92,807 published chromatin interactions from CTCF ChIA-PET in the same cell type. Loops in each chromosome were predicted individually, using models trained from other chromosomes. Genome-wide, we identified 13,360 intra-chromosomal loops. Aggregate peak analysis (APA) shows that there is an enrichment of Hi-C signals in the predicted loop regions (Fig. 2c). Among the predicted loops, 51% have convergent CTCF binding motifs in binding sites, 34% contain CTCF binding sites at a single anchor, and 9% contain tandem motifs (Fig. 2c). 29% of the predicted loops contained promoters at one anchor and enhancers at another, and 41% contained enhancers at both anchors (Fig. 2b). 18% of the predictions contained neither promoters nor enhancers at either anchor.

Next, we trained a model using the 6,395 GM12878 H3K27ac HiChIP loops^37^, which contain more short-range interactions. In total, this model predicted 13,292 loops in the Hi-C matrix. We noticed that the majority of the predictions are the same with the model trained with CTCF: 65% of which exactly matched 64% of predictions from the CTCF model (8,606 of 13,292 and 13,360). When allowing for mismatches of two bins for either anchor, the overlap increases to 69% and 68% (9,109 H3K27ac loops matching 9,135 CTCF loops). The enrichment for promoters and enhancers is also similar to the CTCF ChIA-PET model (Fig. 2b): 29% of the predicted loops are between candidate enhancers and promoters, and ~41% are between enhancers and enhancers.

However, there are differences in the predictions from the two models, and they vividly reflected the difference in the positive training data: 1) we observed a higher percentage of short-range interactions in predictions from the H3K27ac HiChIP model. 53% (7,000/13,292) are short-range (<250kb) while 14% are over 500kb (Fig. 2a). On the contrary, only 36% (4,874/13,360) of predictions by the CTCF model were less than 250kb, while 25% (3,335/13,360) were greater than 500kb (Fig. 2a). This suggests that H3K27ac HiChIP model identifies more short-range interactions, and CTCF ChIA-PET model is better at identifying long-range loops.

Further, we observed a lower CTCF percentage when training with H3K27ac HiChIP, compared with the model trained using CTCF ChIA-PET data (Fig. 2c): 35% of the loops have convergent CTCF binding motifs (vs. 51% in the model trained with CTCF ChIA-PET), 42% contain binding sites at one anchor, and 13% contain no CTCF binding sites. These disparate distributions of CTCF patterns in the prediction are consistent with the patterns in the positive training sets of both models (Supplementary Fig. 3c).

Despite the difference in genomic distance and CTCF motif composition, we observed high validation rates for both prediction sets (Supplementary Fig. 4). 84% (11,151 of 13,292) of loops from the H3K27ac HiChIP model and 83% (11,143 of 13,360) of the CTCF ChIA-PET models can be supported by at least one source, while 61% of HiChIP model and 55% (7,414 of 13,360) of ChIA-PET model can be supported by at least two sources (Supplementary Fig. 4). Considering predictions unique to either model, we found that 66% (2,789 of 4,225) of loops from the CTCF model and 68% (2,824 of 4,183) from the H3K27ac model could be supported by at least one source. At least two sources could support 27% (1,146 loops) of predictions unique to the CTCF model, and 44% (1,827 loops) unique to the H3K27ac model.

Therefore, we decided to use the merged non-redundant predictions with both CTCF ChIA-PET and H3K27ac HiChIP data for GM12878 and all other cell types. Overall, this set of predictions in GM12878 has high recall and validation rates (Fig. 2e and 2f) when compared with four validation sets. Nearly 80% of the Peakachu-predicted loops can be supported by at least one method. At the same time, over 80% of the SMC1 HiChIP interactions can be captured from Peakachu loops at each genomic distance range. The recall rate is lower for CTCF ChIA-PET because that this dataset contains 92,807 loops while we only predicted 17,171 from Hi-C.

### Peakachu reveals a unique set of short-range interactions between enhancers and promoters

We compared Peakachu with HiCCUPS, a popular local enrichment-based method^22^. Peakachu predictions were done on the GM12878 10kb matrix, while HiCCUPS predictions were merged from results on 5kb or 10kb matrices, and an example region is shown in Fig. 2d. First, we observed that 81% (7,639 of 9,448) of HiCCUPS results overlap with our predictions (Supplementary Fig. 3a). Peakachu predicted an additional 10,221 chromatin loops, although the two sets of predictions have similar promoter and enhancer enrichment percentage (Fig. 2b).

We further studied the characteristics of the 10,221 Peakachu unique predictions. 51% (5,162 loops) of them had genomic spans <250kb, suggesting that we identified a unique set of short-range interactions relative to HiCCUPS (Supplementary Fig. 2). 85% (8,726 of 10,221) of the Peakachu-unique loops contained active CTCF binding sites at least one anchor, and 29% (2,974 of 10,221) contain convergent CTCF binding motifs (Supplementary Fig. 2). As for regulatory activity, we found 84% (8,623 of 10,221) of Peakachu-unique loops contained either candidate promoters or enhancers at their anchors, and 31% (3,174 of 10,221) contained candidate promoter-enhancer interactions (Supplementary Fig. 2b). Most importantly, 65% (6,598 of 10,221) of these loops could be validated by at least one ChIA-PET or HiChIP dataset, suggesting that Peakachu identified an additional set of high-confidence biologically meaningful interactions in the genome.

Therefore, we conclude that while searching only for the strongest dot signals on a Hi-C map can reveal a large set of chromatin interactions, a supervised machine-learning approach, especially when trained with H3K27ac HiChIP data, can recover another set of chromatin interactions whose signal patterns might be hard to be identified through local enrichment.

### Estimating false discovery rate (FDR) for Peakachu

In order to estimate the FDR of our model, we applied Peakachu to predict loops for a system previously used to investigate the impact of cohesion loss on loop formations^42^. This system used a modified human colorectal carcinoma cell line HCT-116, with an AID domain tagging to both RAD21 alleles, an indispensable component of the cohesion complex. When treated with auxin, RAD21 in this cell line was effectively destroyed, and loops concomitantly disappeared genome wide due to loss of the essential role of cohesion on loop formation.

We applied our model trained with CTCF ChIA-PET to predict loops in this dataset. Interestingly, Peakachu identified only 19 loops genome-wide from the Hi-C map of auxin-treated cells; in contrast, the same models identified 11,814 loops from the Hi-C map of untreated cells (Supplementary table 2). Given that the sequencing depths are similar between both maps, we can roughly estimate the FDR of Peakachu as ~0.2% (19/11,814).

### Peakachu is robust to sequencing depths from 2 billion to 270 million reads

To test the effect of sequencing depth on the performance of Peakachu, we computationally down-sampled the GM12878 dataset into varying depths (from 100% to 10%) and evaluated its predicted loops. The data was still processed in 10kb resolution at each sequencing depth. First, we observed that lower sequencing depths generally led to reduced number of predicted loops (Fig. 3a). However, the loci of predicted loops are rather stable, using different sequencing depths. For example, there is a large overlap between the predictions that used 10%, 50% and 100% of total sequencing reads (Fig. 3b and 3c). 82% (6,290 of 7,683) of the predicted loops using 10% of the reads are the same as predictions using 2 billion reads.

**Figure 3.**
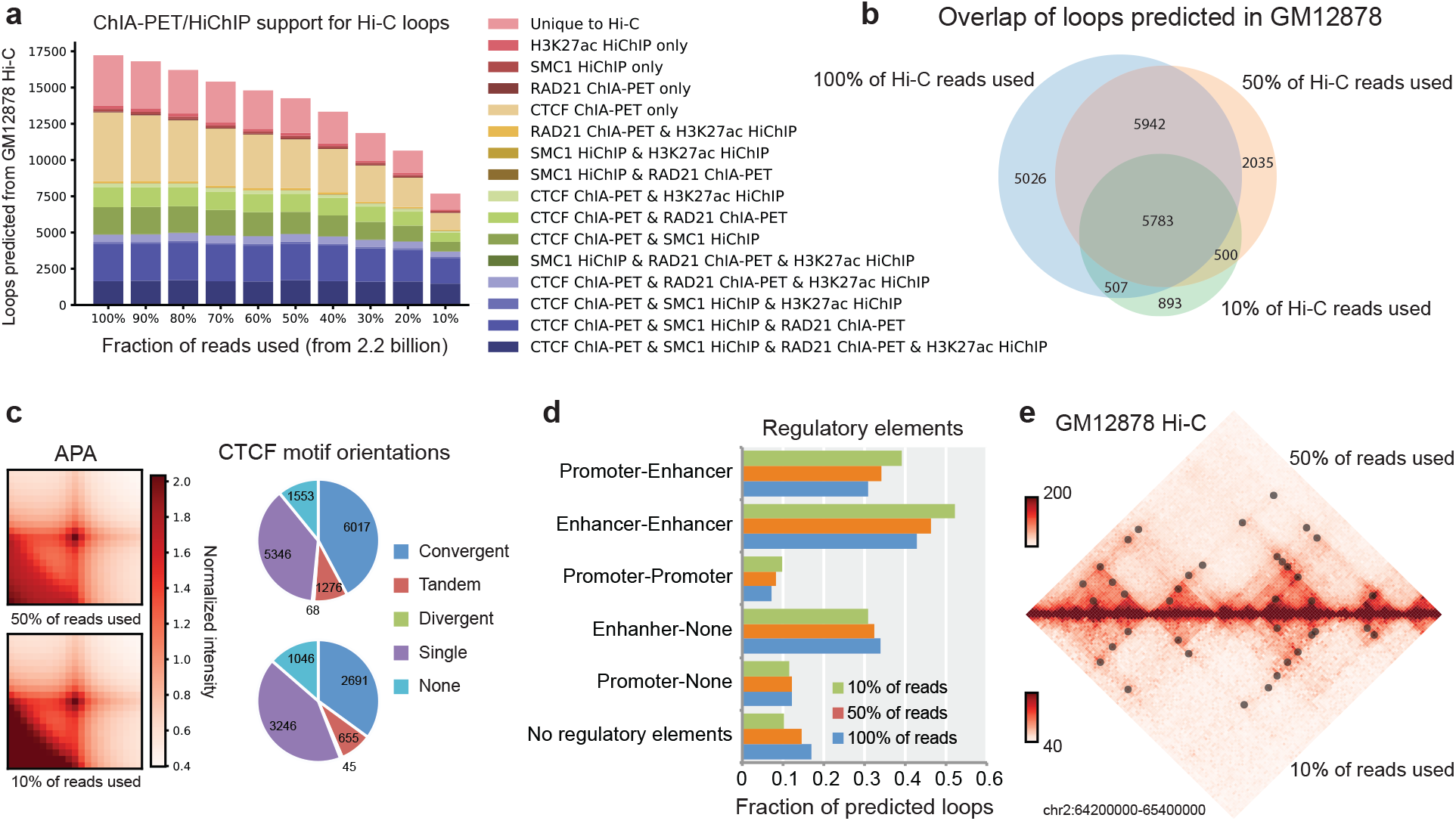
Detection of loops at lower sequencing depths. **a**. Predicted interactions from down-sampled versions of the GM12878 Hi-C map with levels of validation by orthogonal methods. **b**. Concordance of predictions from 100% (blue), 50%, and 10% datasets. **c**. APA profiles and CTCF motif orientations of 50% and 10% loop predictions. Colorbar indicates contact counts normalized by the mean of pixels within their windows (P2M). **d**. Regulatory elements at anchor points of 100%, 50%, and 10% loop predictions. **e**. Visualization of a chromosome 2 locus for Hi-C data and predicted loops using 50% and 10% of sequencing reads.

More importantly, even at 10% down-sample rate, which yields fewer than 300 million read pairs, ~86% (6,582/7,683) of the Peakachu predicted loops can be validated by at least one orthogonal dataset (Fig. 3a). As sequencing depth decreased, validation rates of predicted loops remained similar while their distance distributions tended toward shorter range and retained a majority of loops predicted by HiCCUPS in the same maps (Fig. 3c and 3e, Supplementary Figs. 5 and 6). APA analysis of predicted loops showed enrichment of contact signals compared to surrounding pixels (Supplementary Fig. 7). Further, both CTCF enrichment and ratios of regulatory elements are consistent throughout different sequencing depths (Fig. 3c and 3d). Overall, the Peakachu framework can predict valid interactions with as few as ~270 million sequencing reads in Hi-C data, and predictions for close-range interactions in particular are conserved across sequencing depths.

### Peakachu trained on one cell type accurately capture loops in other cell types

To test whether models trained in one model can be applied in other cell types, we first used the model trained in GM12878 CTCF ChIA-PET to predict loops in the human chronic lymphocytic leukemia cell line K562 (~500 million *cis*-reads) and in mouse embryonic stem cells (1.9 billion *cis*-reads). To match the sequencing depths in K562 and mESC, we used the models trained with 20% and 90% GM12878 Hi-C reads and CTCF ChIA-PET examples (Fig. 4). In K562, we predicted 13,566 chromatin loops: 37% (5,076/13,566) of them contain convergent CTCF binding sites and 41% (5,623/13,566) to have CTCF binding at both anchors, and an additional 45% (6,066 loops) having CTCF binding at one anchor (Fig. 4b). Similarly, we predicted 14,842 loops in mESC: 41% (6,102 of 14,842) of them in convergent orientation and an additional 44% (6,546 of 14,842) with one CTCF anchor (Fig. 4e). Both sets of predictions contained regulatory elements in at least 80% of candidate loops (Fig. 4c and 4f).

**Figure 4.**
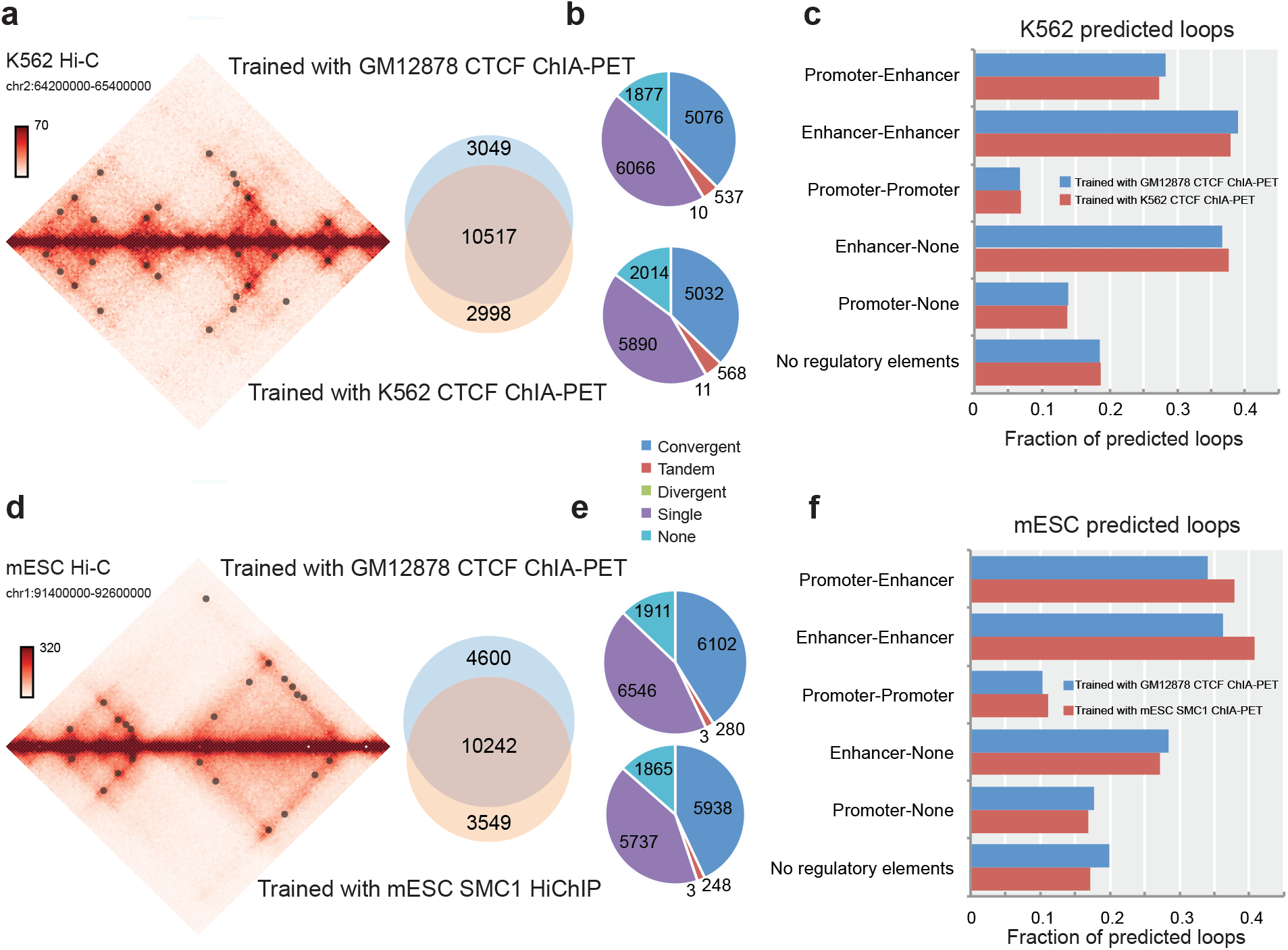
Application in other Hi-C datasets of different tissues and species. **a**. Overlap of loops predicted in K562 Hi-C by Peakachu models trained in either GM12878 or K562 data. Both models were trained with CTCF ChIA-PET examples. **b**. CTCF binding orientations for predicted loops. **c**. Regulatory elements at anchor loci of predicted loops. **d**-**f**. Repeat analysis for loops predicted in mESC Hi-C by Peakachu models trained with either GM12878 CTCF ChIA-PET or mESC SMC1 HiChIP.

Next, we compared predicted loops in K562 from the model trained with GM12878 CTCF ChIA-PET with a model trained with K562 CTCF ChIA-PET. Overall, the predictions are highly similar. A total of 13,566 candidate loops were predicted by the GM12878 model vs 13,515 by K562 model. 78% (10,517/13,566) of GM12878 model and 78% (10,517/13,515) of loops from the K562 model are the same. Their percentage of convergent CTCF binding sites and percentage of enrichment of cis-regulatory elements are consequently similar as well (Fig. 4a and 4c).

For further validate the transferability of Peakachu models, we compared loops predicted in mESC by models trained with either GM12878 CTCF ChIA-PET or mESC SMC1 HiChIP. Again, the total number of predictions was similar, with 14,842 candidate loops from the GM12878 model and 13,791 from the mESC model. Of these, 10,242 were the same, representing 69% of predictions from the GM12878 model and 74% of those from the mESC model (Fig. 4d). While the total overlap was slightly less than the comparable K562 analysis, we found that both sets of mESC predictions had similar distributions for both CTCF binding site orientations (Fig. 4e), and that regulatory elements were slightly more enriched in the model trained with mESC SMC1 ChIA-PET (Fig. 4f).

With K562 and mESC loops serving as a proof-of-concept for transferable GM12878-trained models in other cell types and species, we next predicted interactions in 53 additional Hi-C datasets ranging from 20 million to 3.6 billion *cis*-reads using models trained with down-sampled GM12878 contact maps according to each dataset (Supplementary Table 2 and Supplementary Fig. 10).

### Application of Peakachu framework on DNA SPRITE and Micro-C contact maps

We were interested in the potential of cross-platform comparison enabled by Peakachu. To this end, we tested the performance of Peakachu in Micro-C^9^, a variant of Hi-C protocol capable of higher contact resolutions, and DNA SPRITE^17^, which interrogates chromatin interactions by using a split-pool procedure and assigning a unique sequence barcode for each chromatin contact. We downloaded the H1-ESC Micro-C data from the 4DN web portal (https://data.4dnucleome.org/experiment-set-replicates/4DNES21D8SP8/) and the GM12878 DNA SPRITE data from Quinodoz et al. In both cell lines, CTCF ChIA-PET data are available and were used as positive training sets for Peakachu models trained for Hi-C, Micro-C, and SPRITE (Fig. 5). We used the same parameters and search space for all the analyses.

**Figure 5.**
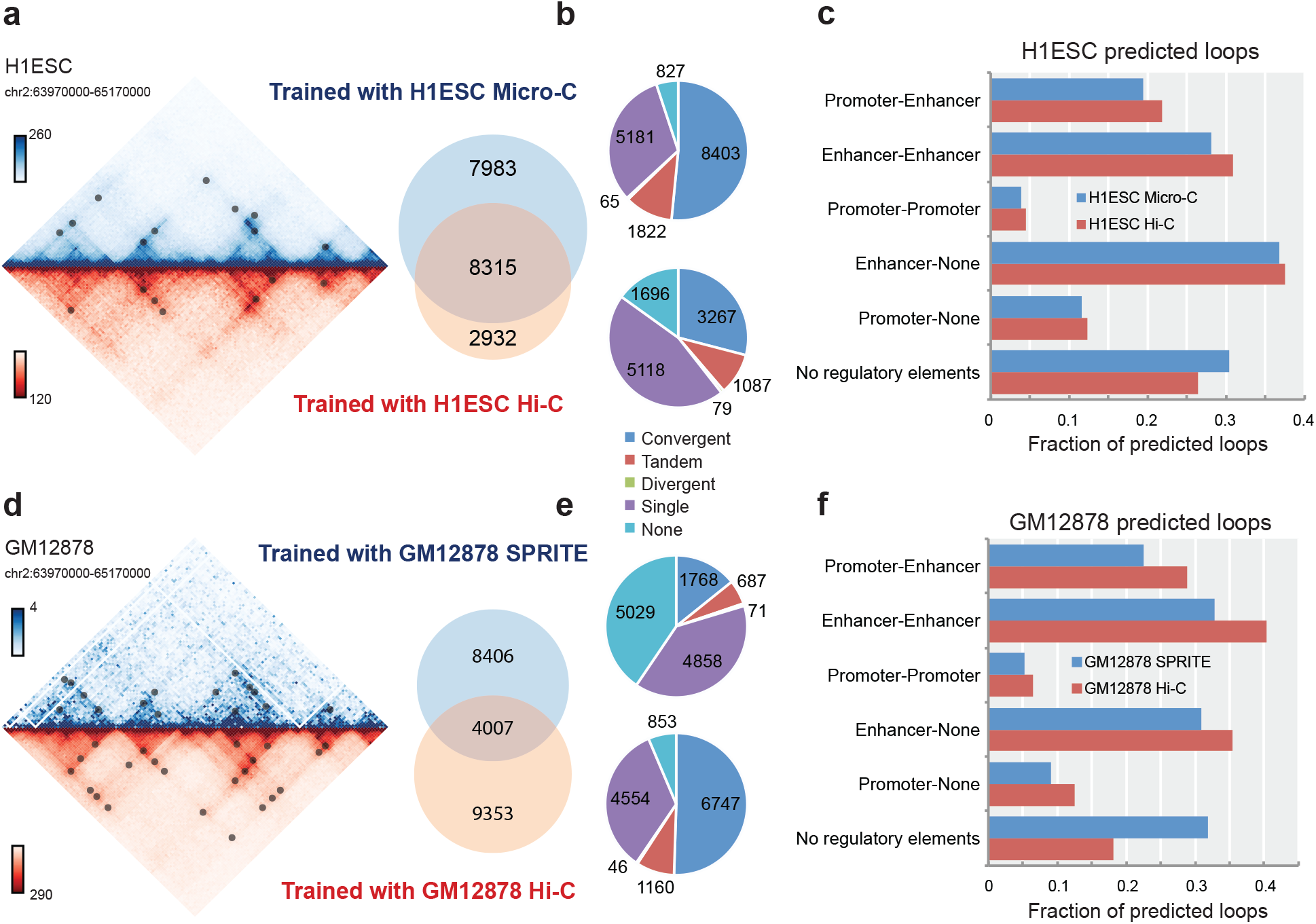
Cross-platform comparison of Peakachu loops in different tissue types. **a**. Overlap of loops predicted in H1ESC by Micro-C and Hi-C, using the same CTCF ChIA-PET training set. **b**. Patterns of CTCF binding site orientations in predicted loops. **c**. Regulatory elements at anchor loci of predicted loops. **d**-**f**. Repeat analysis comparing DNA-SPRITE with Hi-C in GM12878, using the same CTCF ChIA-PET training set.

In Micro-C, we predicted a total of 16,298 loops, which is higher than the loops we predicted in Hi-C data (11,247 loops). This is potentially due to the fact that Micro-C improved the experimental protocol by using a different crosslinker and fragmenting chromatin with higher-resolution enzyme MNase^9^. 74% (8,315/11,247) of the Hi-C loops and 51% of the Micro-C loops are the same (Fig. 5a). Interestingly, the Micro-C predicted loops contain a higher convergent CTCF ratio (52% vs 29%, Fig. 5b) and in general, a higher CTCF binding rate. However, Hi-C predicted loops contain a slightly higher percentage of enhancer-promoter or enhancer-enhancer interactions (Fig. 5c).

The DNA SPRITE dataset for GM12878 contained 135 million *cis*-reads, resulting in a contact map that was quite sparse compared to the Hi-C map comprised by 2.2 billion reads. Despite the disparate depths, Peakachu models trained for SPRITE and Hi-C agreed on 4007 loops in GM12878, representing 32% (4,007/12,413) of SPRITE loops and 30% (4,007/13,360) of Hi-C loops (Fig. 5d). Nearly all SPRITE loops (94%, 11,708 of 12,413) occurred within 250kb, most likely due to the sparsity of the SPRITE contact map. By comparison, 64% of Hi-C loops had genomic spans exceeding 250kb, indicating that SPRITE predictions largely conserved short-range predictions from Hi-C. However, the distributions for CTCF binding orientations and regulatory elements differed greatly between datasets (Fig. 5e and 5f). To further investigate this, we overlapped predictions with CTCF ChIA-PET, RAD21 ChIA-PET, SMC1 HiChIP, and H3K27ac HiChIP datasets and found that 48% (5,990/12,413) of SPRITE loops could be supported by at least one source, vs 84% of Hi-C loops (Supplementary Fig. 9).

Overall, these results support the claim that a general data-driven framework can produce viable decision functions to classify loops in multiple platforms, especially for deeply-sequenced contact maps.

## DISCUSSION

Here we present Peakachu, the first machine learning framework to predict loops from genome-wide contact maps. As we have shown with well-defined training sets from biologically enriched experiments such as ChIA-PET and HiChIP, this totally data-driven approach allows us to detect high-quality loop interactions in various platforms and sequencing depths. Importantly, the model is transferable among different datasets from the same platform. With the model trained in GM12878, we systematically performed loop predictions in 55 additional Hi-C datasets, the sequencing depths ranging from 25 million to 3.6 billion *cis*-reads.

An open question in the field of 3D genomics is how to define loops. There already exist many methods for loop detection. However, to the best of our knowledge, all of them are based on searching for statistically enriched interactions against a global or local background. Due to different choice of statistical model and different background definitions, these methods can yield disparate results even for the same dataset. Our motivation of this work is to minimize artificial definition of loops and learn inherent patterns from data itself. We treated this task as a binary classification problem and selected the widely used random forest model as the learner. We mainly tested two sets of interactions for model training: one is CTCF ChIA-PET, enriched with long-range structural interactions; the other is H3K27ac HiChIP, enriched with short-range regulatory interactions. Strikingly, the loops predicted by different model recapitulated the main features of the corresponding training sets. One potential extension of this framework will be training with interactions from various orthogonal experiments, even with imaging data such as HiFISH, to reveal different subsets of chromatin loops throughout the genome.

Because most cell lines or tissues only have low-sequencing depth Hi-C data available, we investigated the impact of sequencing depth on our framework by down-sampling the deeply sequenced GM12878 dataset and repeating training and predicting on the series of contact matrices with varying depths. Although the sensitivity reduced along the decreasing sequencing depths, most loop predictions remained stable even in the 10% down-sampled datasets (less than 300 million cis- reads). After further proving the transferability of Peakachu models trained in GM12878 in K562 and mESC, we predicted loops in 53 additional cell lines or tissues. We found that even in datasets with ~25 million cis-reads, Peakachu still successfully detected ~4,500 loops with distinct loop patterns according to APA profiles (Supplementary Table 2 and Supplementary Fig. 10). We have made the predicted loops in all the 56 Hi-C datasets available in our 3D genome browser, which will serve as a resource for future studies that utilize the same cells in the list.

The number of techniques for chromosome conformation study continues to grow, and several such as GAM and DNA SPRITE still lack of dedicated algorithms for chromatin loop detection. Here we show the generalizability of our framework by testing its performance on contact matrices from GM12878 DNA SPRITE and human embryonic Micro-C experiments. While loops predicted from Micro-C and Hi-C are similar, Peakachu detected quite a few specific biologically meaningful loops in DNA SPRITE compared with Hi-C. This could be explained by disparate experimental pipelines adopted by these two techniques. In future studies, we will apply Peakachu framework in all available platforms to investigate advantages and pitfalls of each techniques in loop detecting and unveil the complete picture of loop-level structure within chromosomes.

## METHODS

Fitting a Peakachu model requires two components: a Hi-C matrix binned to 10kb and a ChIA-PET interaction list. For every interaction in the list, a corresponding 11×11 window centered at the interaction is collected from the Hi-C matrix. The ratio of the center pixel to the lower left quadrant (P2LL) of this window is used as an indicator variable prior to training, and the minimum P2LL for the positive class is set to 0.1. In other words, samples from the input training list are rejected if their Hi-C value is less than 10% than the average value within the loop. After collecting the positive class, a comparable number of windows with random coordinates and nonzero centers are collected to define a negative class.

Each sample is decomposed into a vector of 243 features. 121 of these represent the values of each pixel in an 11×11 window. Another 121 represent the relative ranks of each pixel within the window. A final variable, P2LL, is appended to each vector of features. Using scikit-learn, the training set is then passed to a 3-fold cross-validation loop to fit a random forest of 100 decision trees; each tree is trained on a random combination of 15-20 features, and the hyperparameters tested include splitting criterion (entropy or gini), maximum tree depth, and class weights. Matthew’s Correlation Coefficient is used as the primary metric for selecting the best model. Training with a GM12878 Hi-C matrix with a list of 76,667 CTCF interactions and 78,069 random loci requires 12 minutes of runtime and a peak RAM usage of 4Gb.

Trained models can be applied to contact maps of the same platform and similar depth as the training matrix. Peakachu defines feature vectors for all nonzero pixels within a given genomic span, and scores each using the predict_proba method provided by the scikit-learn library. Usually, highly scored pixels are found grouped together, and only one representative pixel is reported from each cluster. To select representative pixels, we developed a greedy algorithm entailing two steps: first, define 1D loop anchor regions enriched for highly scored (P>0.9) pixels; then run DBSCAN between any two connected anchor regions. The identification of the loop-enriched anchors was performed by counting the candidate pixels and finding peaks along the chromosomes. Specifically, we applied the find_peaks and peak_widths functions from Python’s Scipy package to locate the peak summits and estimate the peak widths respectively.

When applying HiCCUPS on 10kb Hi-C maps, we noticed little variance in results for different parameter settings. We therefore used default settings to predict loops and separately considered the published HiCCUPS looplist containing predictions from 10kb and 5kb matrices.

## Acknowledgements

F.Y. acknowledged the support from National Institutes of Health (NIH) [1R35GM124820, R01HG009906, U01CA200060 and R24DK106766]. We are thankful to members of the Yue lab for the discussions and suggestions.

## Author contributions

F.Y. designed and supervised the project. T.S. and X.W. implemented the PEAKACHU software. F.S., B.Z., S.M.W., C.K. facilitated with the data analysis. T.S., X.W., and F.Y wrote the manuscript with input from all the authors.

## Data availability

Source code is publicly available at available in the GitHub repository (github.com/tariks/peakachu). Predictions in 56 cell/tissue types can be downloaded in the 3D Genome Browser (http://3dgenome.org).

---

**Supplementary Figure 1 | Venn diagram of ChIA-PET and HiChIP results in GM12878**. Overlaps were computed with the bedtools pairtopair command with parameters -slop 15000 – type both. In overlap regions, the contribution of each dataset is coded by color and order of the legend.

**Supplementary Figure 2 | Analysis of 10,221 predicted loops unique to Peakachu with respect to HiCCUPS**. **a**. pie charts for active CTCF motif orientations and distance distrubutions of predicted loops. **b**. Non-mutually-exclusive bars for regulatory elements at anchor loci.

**Supplementary Figure 3 | Comparison of Peakachu models trained with CTCF ChIA-PET or H3K27ac HiChIP**. **a**. Venn diagram of loop prediction in GM12878 from two Peakachu models and HiCCUPS. Overlaps defined by loop predictions within 3 bins of genomic distance. **b**. Enriched motifs in predicted anchor regions. **c**. CTCF binding patterns of the training sets for both models.

**Supplementary Figure 4 | Recall and precision of predictions made by Peakachu models trained with different inputs in GM12878**. **a**-**c**. Interactions from orthogonal datasets recaptured from Hi-C predictions by two Peakachu models. **d**-**f**. Ratio of loops with orthogonal support among 4,225 unique predictions from the CTCF model, 4,183 unique predictions from the H3K27ac model, and 9,135 loops predicted by both models.

**Supplementary Figure 5 | Performance of HiCCUPS and concordance with Peakachu for downsampled GM12878 contact maps**. **a**. Overlap of HiCCUPS interactions with ChIA-PET and HiChIP datasets for predictions in GM12878 at different read depths. **b**. Overlap with Peakachu loops predicted from the same contact maps are displayed as Venn diagrams.

**Supplementary Figure 6 | Genomic distance distributions for loops predicted in GM12878 Hi-C by Peakachu and HiCCUPS at various read depths**. The final panel displays distance distributions of orthogonal datasets in GM12878.

**Supplementary Figure 7 | APA profiles of loops predicted in down-sampled datasets**. Non-redundant loops were merged from predictions by CTCF-trained and H3K27ac-trained models.

**Supplementary Figure 8 | Recall and Precision values for loops predicted from Hi-C data**. Loops were predicted by either natively trained models or transferred models from GM12878 training sets.

**Supplementary Figure 9 | Validation and recapture rates for loops predicted in Micro-C and DNA SPRITE datasets**. Loops were predicted using models trained with CTCF ChIA-PET examples, and evaluated using interactions from orthogonal experiments as validation sets.

**Supplementary Figure 10 | APA profiles of loops predicted in 56 Hi-C datasets.** Predictions were made by models trained on CTCF ChIA-PET examples in GM12878 maps of similar depths.

**Supplementary table 1.** Collected genome-wide contact datasets for GM12878.

| Dataset | Publication | Reads | Contacts |
| --- | --- | --- | --- |
| GM12878 Hi-C | Rao..Aiden Cell 2014 | 3,587,190,419 | 9,448 |
| CTCF ChIA-PET | Tang..Ruan Cell 2015 | 51,103,494 | 92,807 |
| RAD21 ChIA-PET | Heidari..Snyder 2014 | 225,434,403 | 20,887 |
| HiChIP SMC1 | Mumbach..Chang<br>Nature Met. 2016 | 634,461,964 | 10,254 |
| HiChIP H3K27ac | Mumbach..Chang<br>Nature Gen. 2017 | 332,279,257 (B1)<br>299,478,736 (B2) | 6,395 |

**Supplementary table 2** (excel sheet) Predicted loop counts in 56 Hi-C datasets.

