## Supplementary Figures for "A supervised learning framework for chromatin loop detection in genome-wide contact maps"

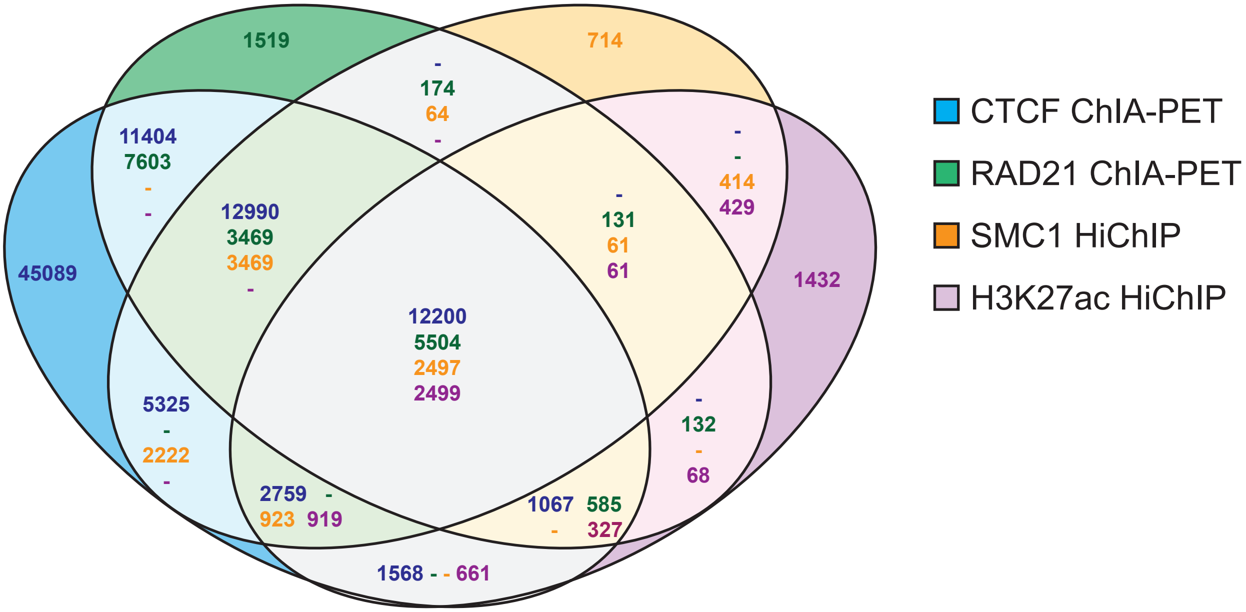


**Supplementary Figure 1. Venn diagram of ChIA-PET and HiChIP results in GM12878**. Overlaps were computed with the bedtools pairtopair command with parameters –slop 15000 –type both. In overlap regions, the contribution of each dataset is coded by color and order of the legend.

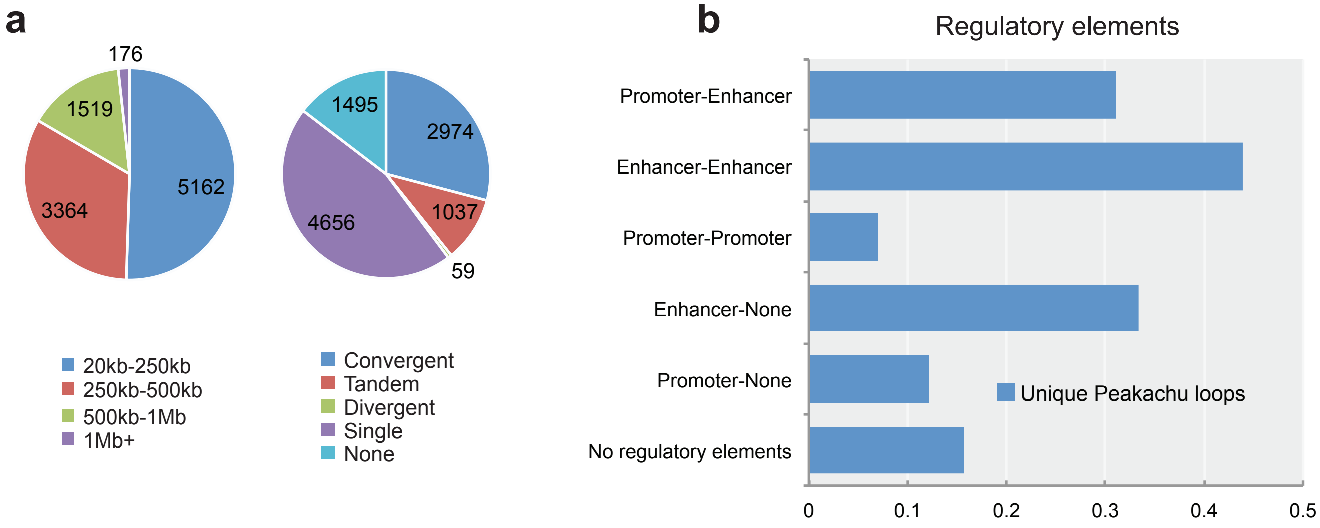


**Supplementary Figure 2. Analysis of 10,221 predicted loops unique to Peakachu with respect to HiCCUPS**. **a.** pie charts for active CTCF motif orientations and distance distrubutions of predicted loops. **b.** Non-mutually-exclusive bars for regulatory elements at anchor loci.

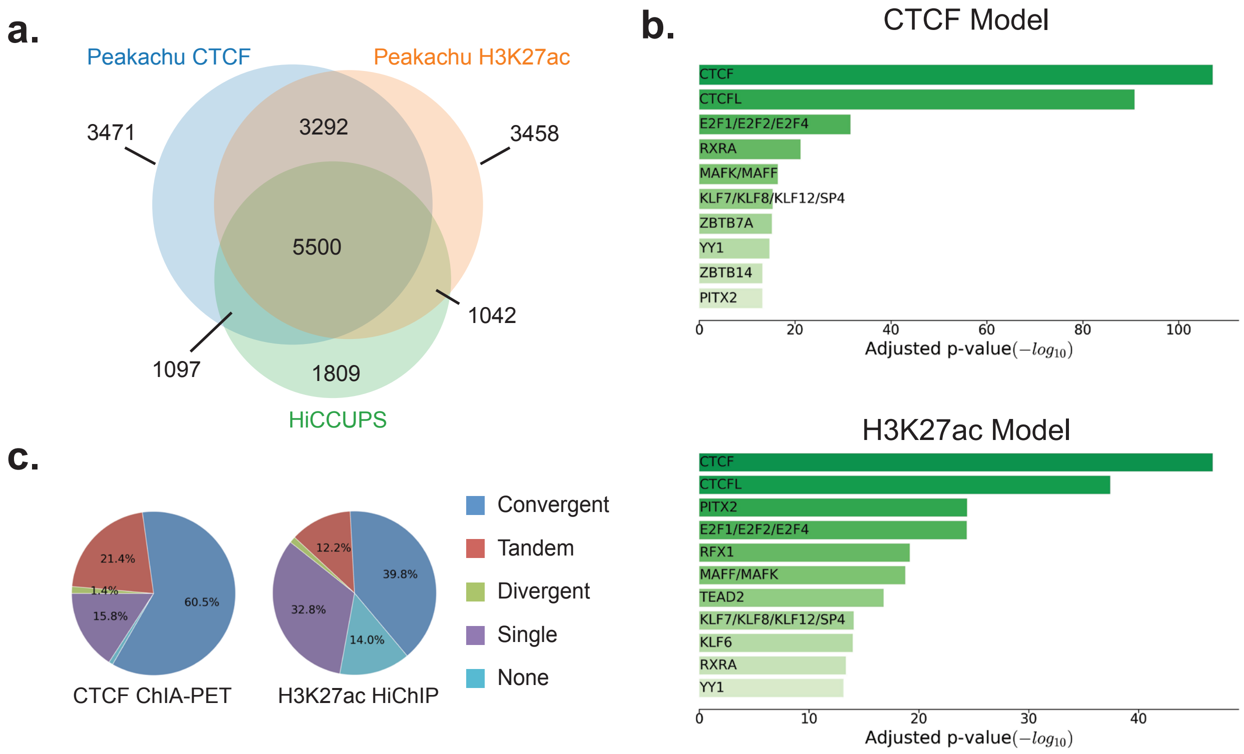


**Supplementary Figure 3. Comparison of Peakachu models trained with CTCF ChIA-PET or H3K27ac HiChIP. a.** Venn diagram of loop prediction in GM12878 from two Peakachu models and HiCCUPS. Overlaps defined by loop predictions within 3 bins of genomic distance. **b.** Enriched motifs in predicted anchor regions. **c.** CTCF binding patterns of the training sets for both models.

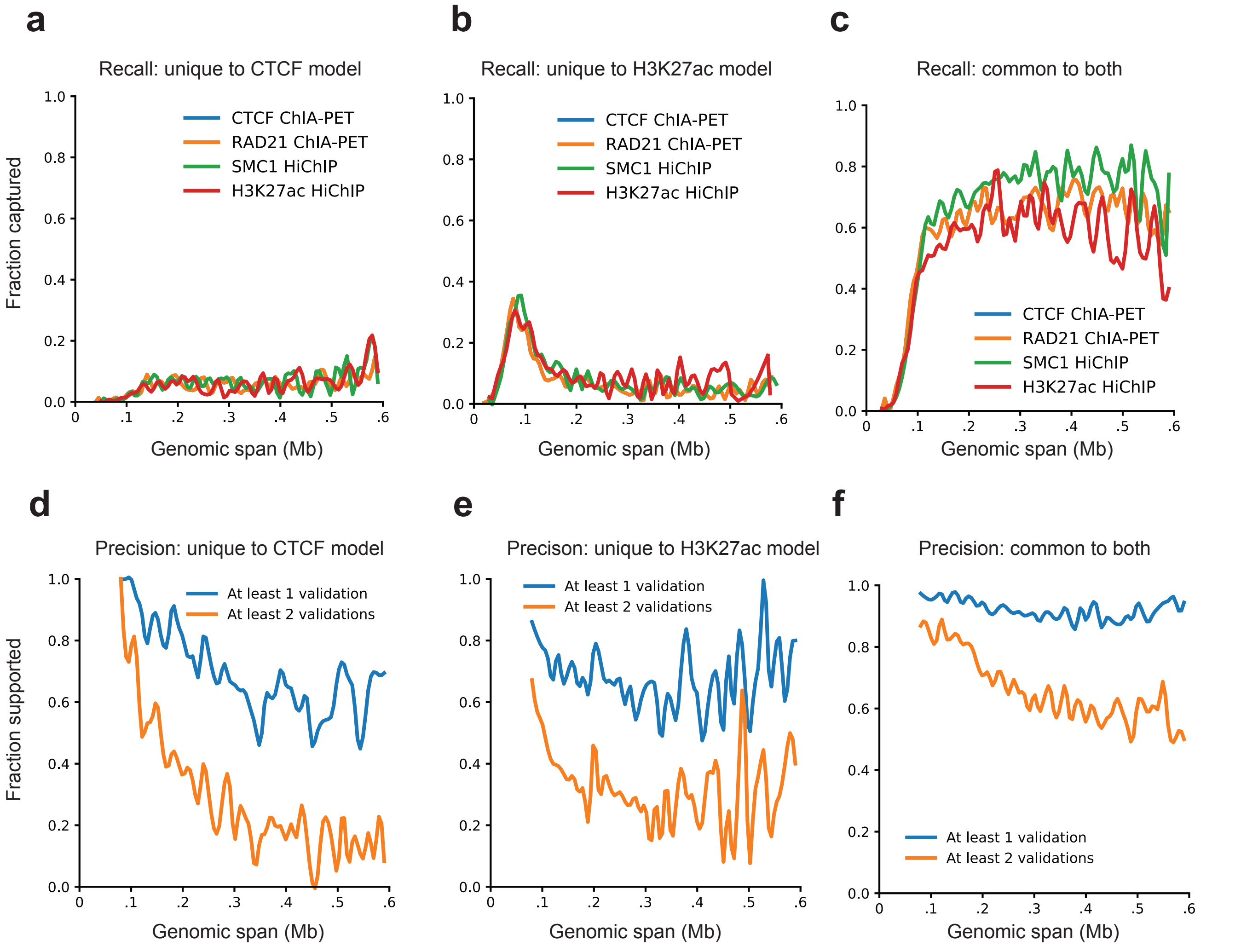


**Supplementary Figure 4. Recall and precision of predictions made by Peakachu models trained with different inputs in GM12878. a-c.** Interactions from orthogonal datasets recaptured from Hi-C predictions by two Peakachu models. **d-f.** Ratio of loops with orthogonal support among 4,225 unique predictions from the CTCF model, 4,183 unique predictions from the H3K27ac model, and 9,135 loops predicted by both models.

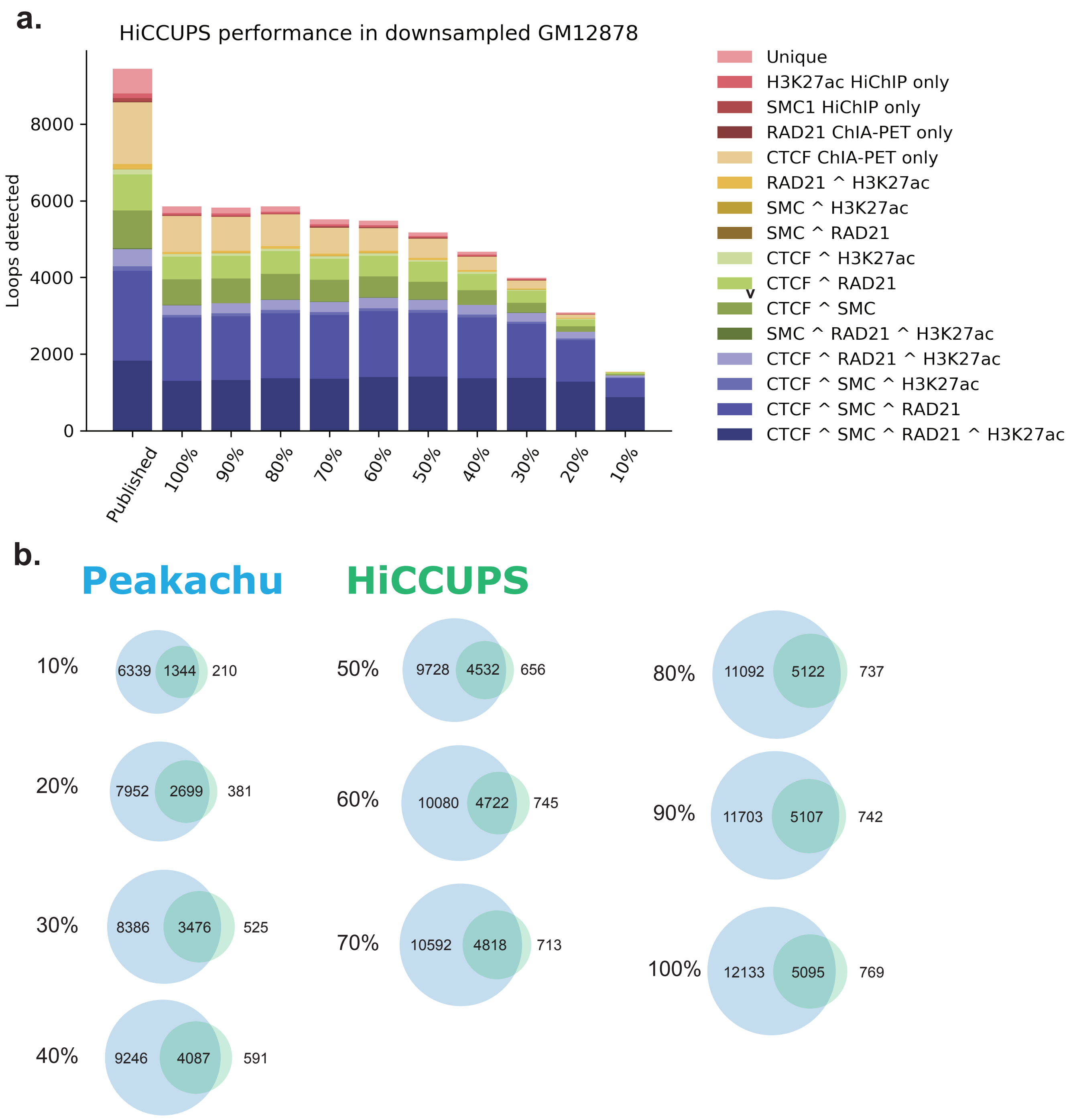


**Supplementary Figure 5. Performance of HiCCUPS and concordance with Peakachu for downsampled GM12878 contact maps**. **a.** Overlap of HiCCUPS interactions with ChIA-PET and HiChIP datasets for predictions in GM12878 at different read depths. **b.** Overlap with Peakachu loops predicted from the same contact maps are displayed as Venn diagrams.

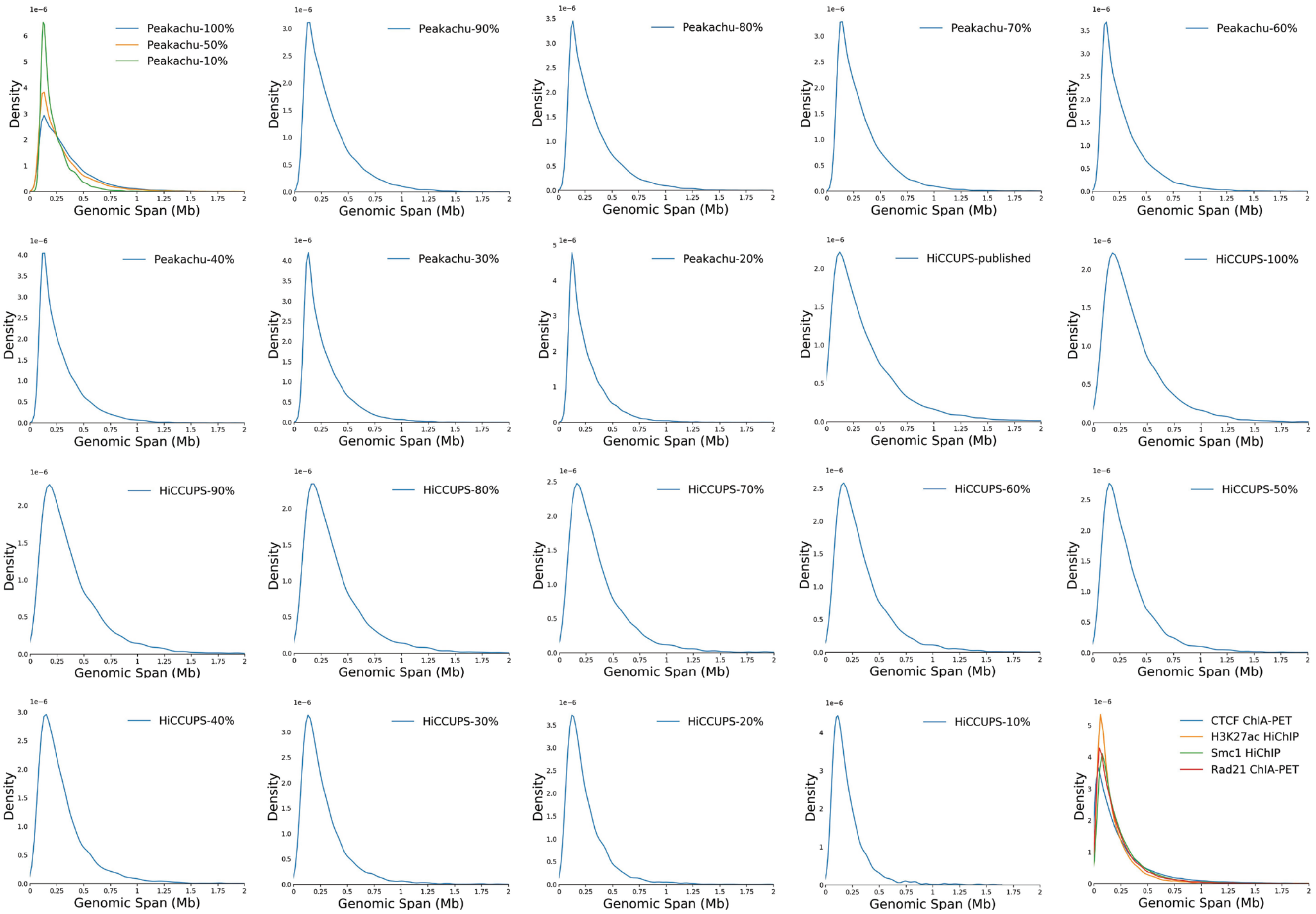


**Supplementary Figure 6. Genomic distance distributions for loops predicted in GM12878 Hi-C by Peakachu and HiCCUPS at various read depths**. The final panel displays distance distributions of orthogonal datasets in GM12878.

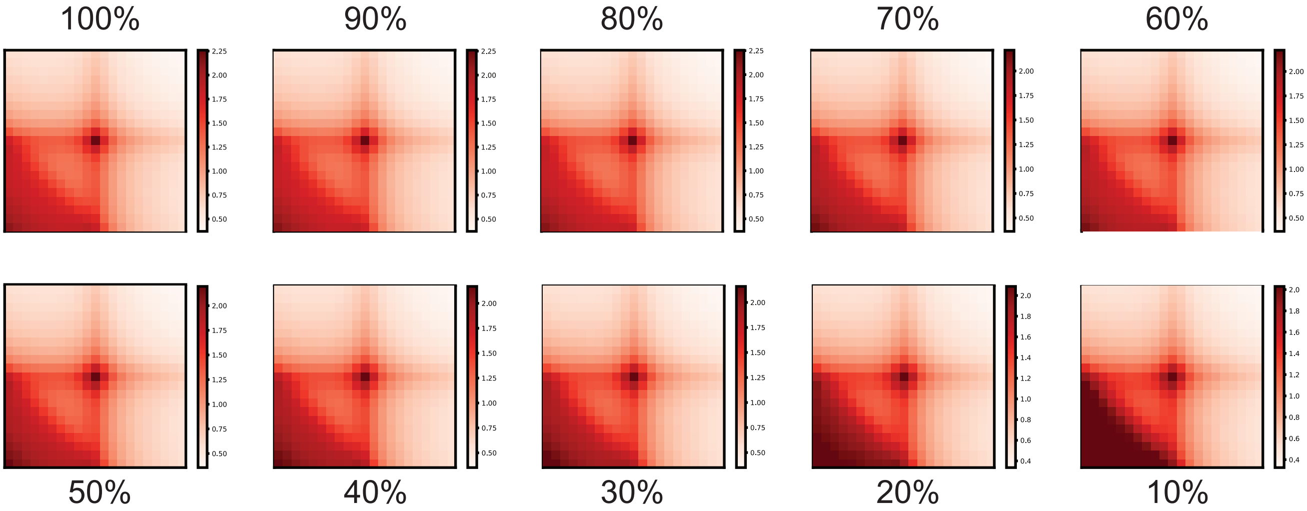


**Supplementary Figure 7. APA profiles of loops predicted in down-sampled datasets**. Non-redundant loops were merged from predictions by CTCF-trained and H3K27ac-trained models.

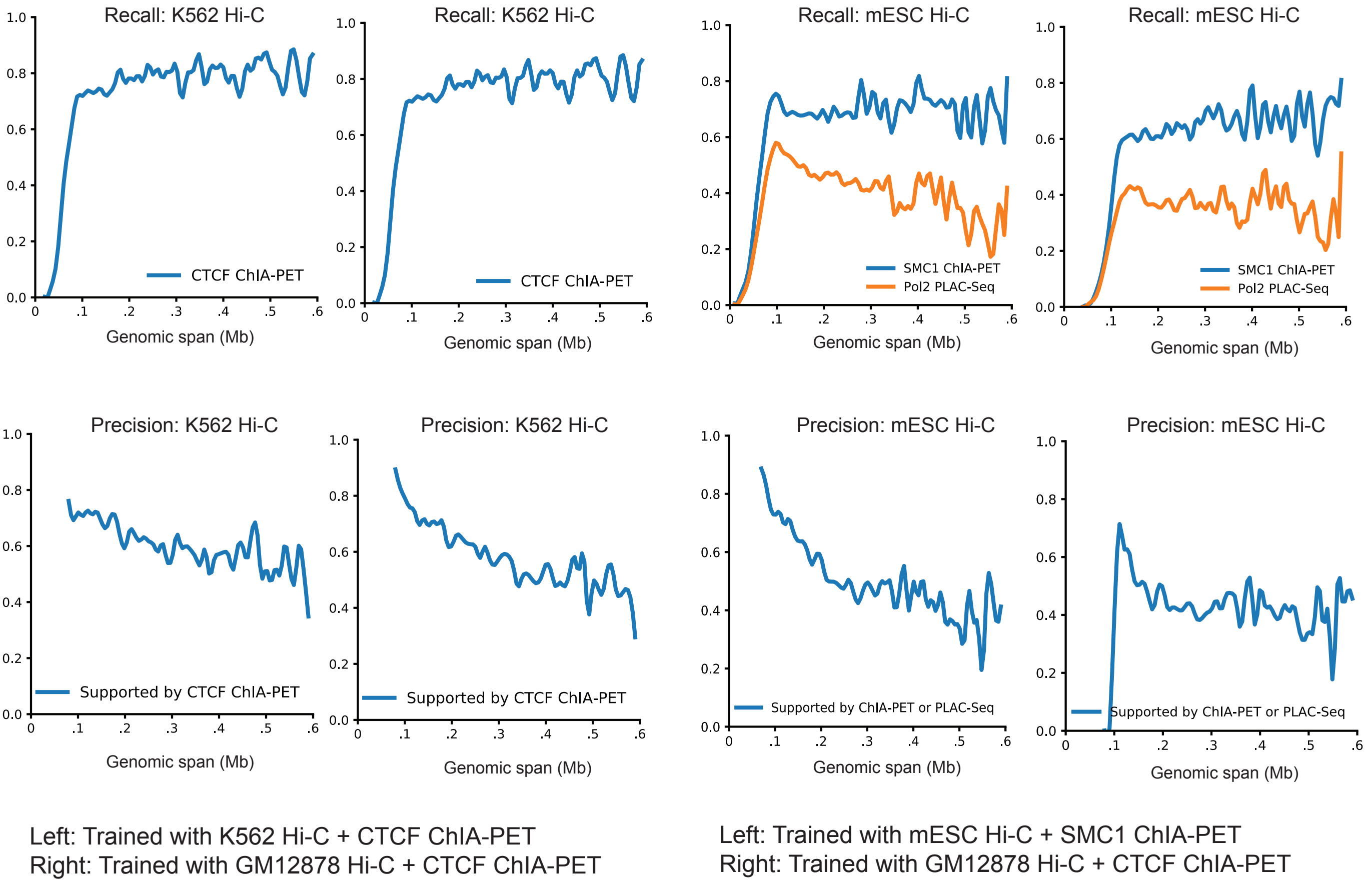


**Supplementary Figure 8. Recall and Precision values for loops predicted from Hi-C data.** Loops were predicted by either natively trained models or transferred models from GM12878 training sets.

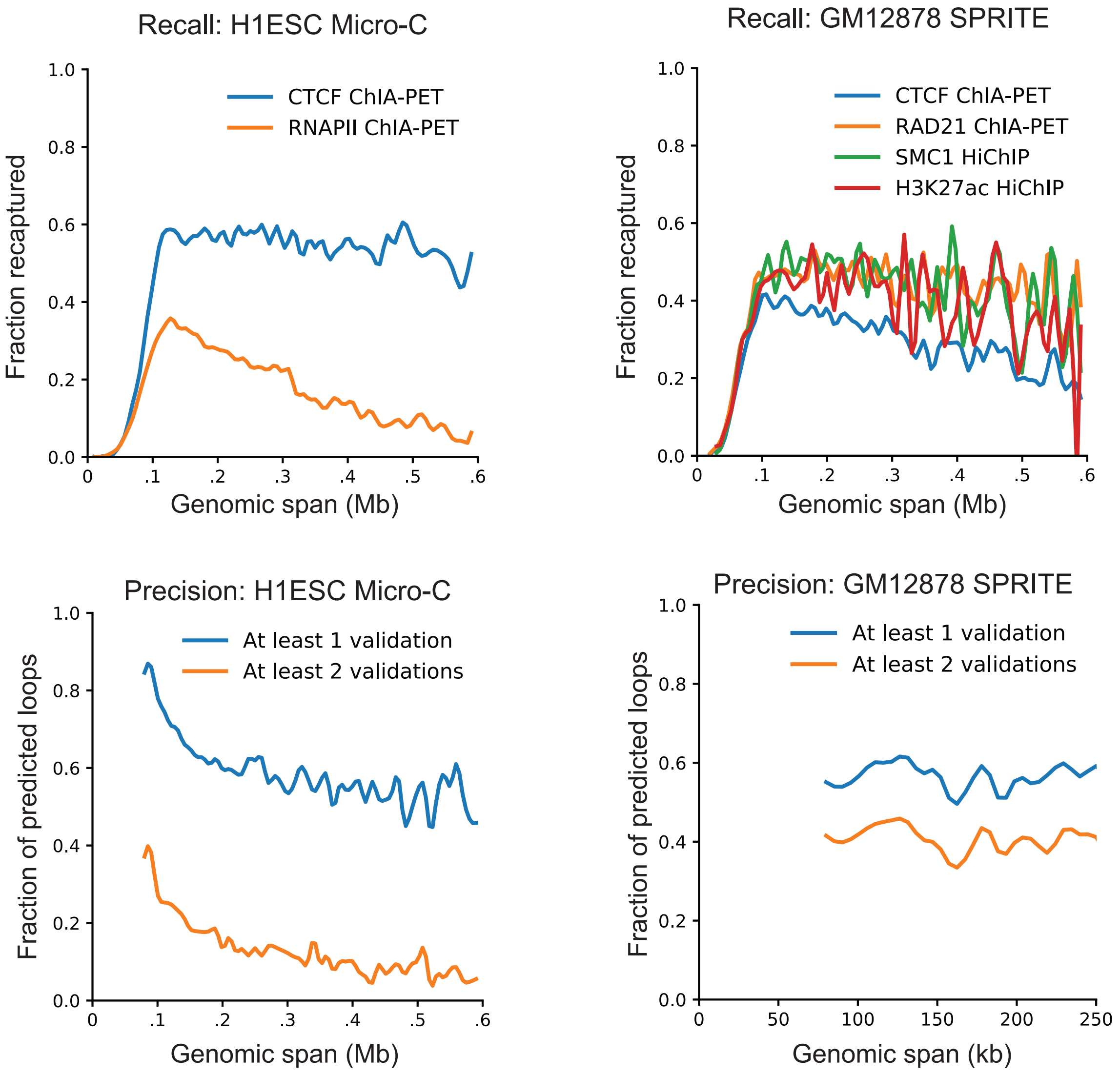


**Supplementary Figure 9. Validation and recapture rates for loops predicted in Micro-C and DNA SPRITE datasets**. Loops were predicted using models trained with CTCF ChIA-PET examples, and evaluated using interactions from orthogonal experiments as validation sets.

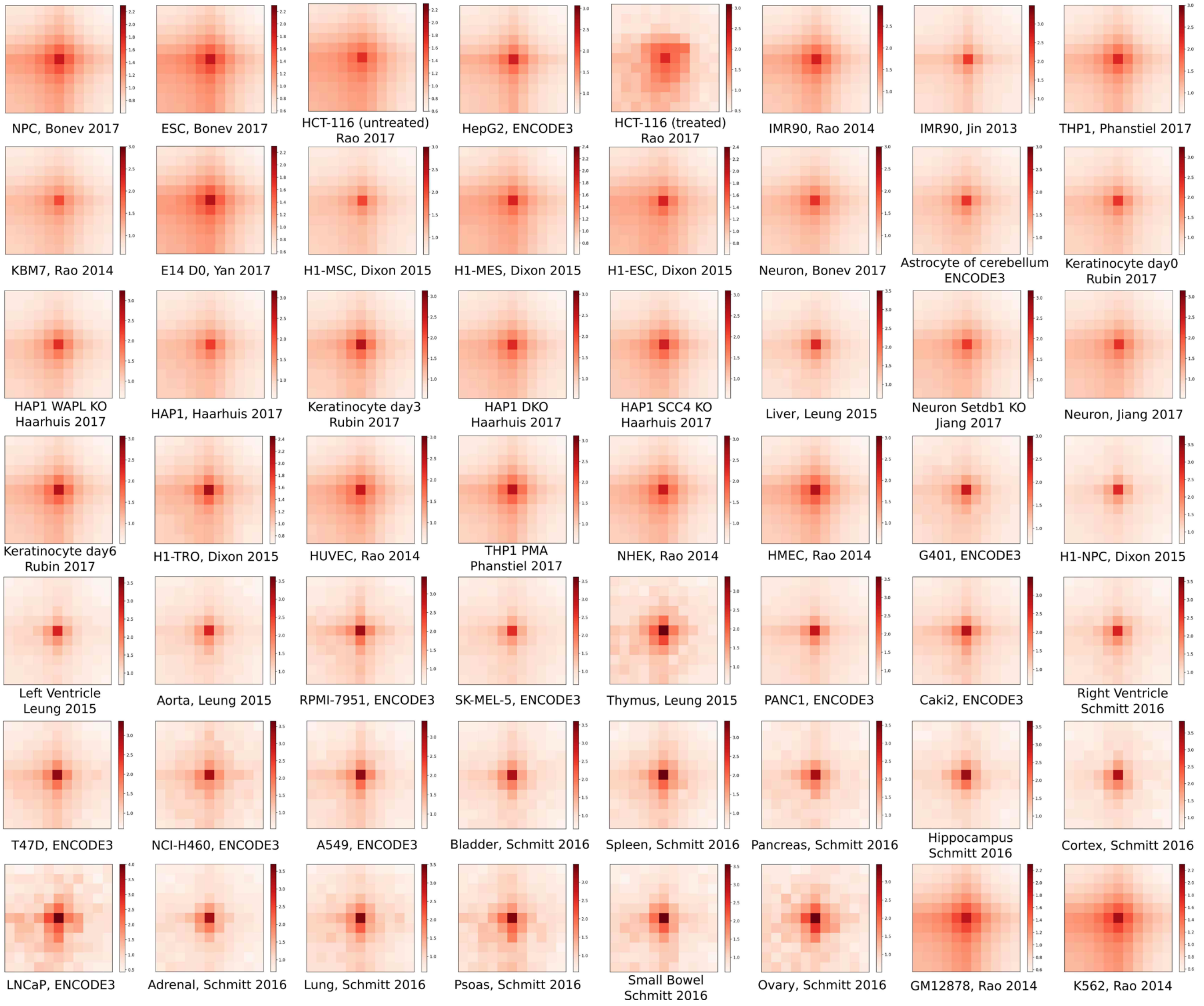


**Supplementary Figure 10. APA profiles of loops predicted in 56 Hi-C datasets.** Predictions were made by models trained on CTCF ChIA-PET examples in GM12878 maps of similar depths.
